# PROTON MOTIVE FORCE INHIBITORS ARE DETRIMENTAL TO METHICILLIN-RESISTANT *STAPHYLOCOCCUS AUREUS* PERSISTER CELLS

**DOI:** 10.1101/2022.05.24.493181

**Authors:** Sayed Golam Mohiuddin, Sreyashi Ghosh, Pouria Kavousi, Mehmet A. Orman

## Abstract

Methicillin-resistant *Staphylococcus aureus* (MRSA) strains are resistant to conventional antibiotics. These pathogens can form persister cells, which are transiently tolerant to bactericidal antibiotics, making them extremely dangerous. Previous studies have shown the effectiveness of proton motive force (PMF) inhibitors at killing bacterial cells; however, whether these agents can launch a new treatment strategy to eliminate persister cells mandates further investigation. Here, using known PMF inhibitors and two different MRSA isolates, we showed that antipersister potency of PMF inhibitors seemed to correlate with their ability to disrupt PMF and permeabilize cell membranes. By screening a small chemical library to verify this correlation, we identified a subset of chemicals (including nordihydroguaiaretic acid, gossypol, trifluoperazine, and amitriptyline) that strongly disrupted PMF in MRSA cells by dissipating either the transmembrane electric potential (ΔΨ) or the proton gradient (ΔpH). These drugs robustly permeabilized cell membranes and reduced persister levels below the limit of detection. Overall, our study further highlights the importance of cellular PMF as a target for designing new antipersister therapeutics.

## INTRODUCTION

The discovery of antibiotics in the 1940s was one of the most significant breakthroughs in therapeutic medicine. However, the medicinal potency of these lifesaving drugs has been drastically reduced by the emergence of new antibiotic-resistant mutant bacterial strains. The continuous evolution of pathogens to develop resistance against antibiotics, together with the decreased rate of antibiotic discovery, might eventually cause serious public health problems, as epidemics associated with resistant pathogens may be imminent. Bacterial persisters, phenotypic variants that are transiently tolerant to high concentrations of antibiotics^1,2^, exacerbate the problem, as they may form a reservoir for the emergence of antibiotic-resistant mutant strains^3^. Bacterial persistence is a non-heritable, reversible, antibiotic-tolerant state that can be triggered by stochastic and/or deterministic factors^4–7^. Persisters are the leading cause of the propensity of biofilm infections to relapse^8^.

*Staphylococcus aureus* is an opportunistic Gram-positive bacterial pathogen that colonizes human skin and mucous membranes, causing chronic, recurrent infections, including wound infections, bacteremia, and biofilm infections^9,10^. Methicillin, a narrow-spectrum beta-lactam antibiotic, was introduced in the late 1950s to treat infections caused by penicillin-resistant *S. aureus*^11^. Unfortunately, accession of the methicillin-resistance gene, *mecA*, encoding an alternative penicillin-binding protein, makes *S. aureus* infections extremely difficult to treat^12^. Methicillin-resistant *S. aureus* (MRSA) emerged as a major hypervirulent pathogen that causes severe healthcare-acquired infections, such as surgical site infections, hospital-acquired pneumonia, catheter-associated urinary tract infections, central line–associated bloodstream infections, and ventilator-associated pneumonia^13^. Almost 19,000 people die annually as a consequence of MRSA infections in the United States (US) alone^14^. Approximately 20% of patients in the US contract at least one nosocomial infection while undergoing surgery, which adds $5–10 billion in costs to the US healthcare system^15,16^.

Quorum sensing, reactive oxygen species, the stringent response, inactivation of the tricarboxylic acid cycle, and the SOS response have been implicated in persister formation in *S. aureus*^17–21^. Reducing intracellular ATP formation with arsenate treatment can induce persistence in *S. aureus*^22^. However, intracellular *S. aureus* persisters isolated from human macrophages are metabolically active and display an altered transcriptomic profile^17^, suggesting that the correlation between high persistence and ATP reduction is not universal. Conventional antibiotics, such as gentamicin (a protein synthesis inhibitor) and ciprofloxacin (a DNA synthesis inhibitor), fail to eliminate MRSA persisters^21,23–25^. Some MRSA strains have already acquired resistance against vancomycin (a cell wall biosynthesis inhibitor)^26^.

The cell membrane is an essential cellular component and might be a good target for novel antipersister therapeutics^27^. The bacterial proton motive force (PMF) maintains the electrochemical proton gradient across the cell membrane, an essential component of ATP synthesis^27,28^. The electric potential (ΔΨ) and the transmembrane proton gradient (ΔpH) are the two components of PMF. Cells can compensate for the dissipation of one component by enhancing the other to maintain the necessary level of PMF^29^. A number of chemicals disrupt PMF of *S. aureus* by dissipating either ΔΨ or ΔpH^30–32^. Halicin is a potential broad-spectrum antibacterial molecule that selectively dissipates ΔpH^31^. The small molecule JD1 disrupts ΔΨ, kills MRSA cells, and significantly reduces biofilm formation^32^. Bedaquiline, SQ109, pyrazinamide, clofazimine, nitazoxanide, and 2-aminoimidazoles are also potent PMF inhibitors in gram-positive bacteria^30,33^. PMF inhibitors can also permeabilize the membranes of metabolically active cells through interactions with phospholipids or membrane-bound proteins^34–36^. Polymyxin B, a well-known inhibitor of ΔΨ, perturbs the cell membranes of bacteria by binding lipopolysaccharides^27,37^.

Although the effectiveness of PMF inhibitors against bacterial cells has been highlighted in prior studies^27,31,34,37–41^, whether PMF inhibitors can be used as an antipersister therapeutic strategy necessitates further investigation. In this study, we sought to determine if disrupting PMF of MRSA persisters can be detrimental for these cells. Because electron transport chain (ETC) complexes are highly conserved across species, we used a small library of 22 chemical compounds that inhibit various mitochondrial ETC complexes and identified several drugs (nordihydroguaiaretic acid, gossypol, trifluoperazine, and amitriptyline) that disrupted PMF in MRSA strains by dissipating either ΔpH or ΔΨ. Although most of these chemicals drastically reduced MRSA survival compared with conventional antibiotics, our subsequent analysis verified that the extent of PMF disruption and membrane permeabilization is a key factor determining the treatment outcome.

## RESULTS

### PMF inhibitors can effectively kill MRSA strains

First, we tested the effectiveness of known PMF inhibitors, such as polymyxin B, thioridazine, and carbonyl cyanide m-chlorophenyl hydrazone (CCCP), on MRSA persistence in two isolates: MRSA BAA-41 and MRSA 700699. Polymyxin B is a cationic peptide that electrostatically binds the negatively charged moieties of lipopolysaccharides, disrupting ΔΨ and permeabilizing the cell membrane^42^. Thioridazine, an antipsychotic drug, disrupts ΔΨ in gram-positive bacteria, potentially by blocking NADH:quinone oxidoreductase II (NDH-II)^36,43^. CCCP is a protonophore that transports hydrogen ions across the cell membrane, subsequently reducing ATP production and disrupting PMF^44^.

Dissipation of either ΔΨ or ΔpH by inhibitors may cause the eventual collapse of the bacterial cellular PMF and disrupt membrane integrity^27^. To assess the effects of PMF inhibitors on membrane permeability, strains MRSA BAA-41 and MRSA 700699 were grown to an optical density at 600 nm (OD_600_) of ∼0.1 in Mueller–Hinton broth in test tubes (**Supplementary Fig. S1**); treated with polymyxin B, thioridazine, or CCCP at 5× or 10× the minimum inhibitory concentration (MIC) for 1 hour (**Supplementary Table S1A**); and then stained with propidium iodide (PI). PI is a membrane-impermeant DNA- and RNA-binding dye that can only stain nucleic acids of cells with compromised membranes. Flow cytometric analysis of PI-stained cells revealed that polymyxin B at 5× and 10× MIC permeabilized more than 80% of MRSA BAA-41 cells (**Fig. 1A** and **Supplementary Table S2A**) but less than 80% of MRSA 700699 cells (**Fig. 2A** and **Supplementary Table S2A**). Although robust membrane permeabilization was not observed after CCCP treatment at the indicated concentrations in either strain (**Fig. 1A, Fig. 2A**, and **Supplementary Table S2A**), thioridazine treatment at 5× and 10× MIC permeabilized more than 90% of cells of both strains (**Fig. 1A, Fig. 2A**, and **Supplementary Table S2A**).

**Fig. 1.**
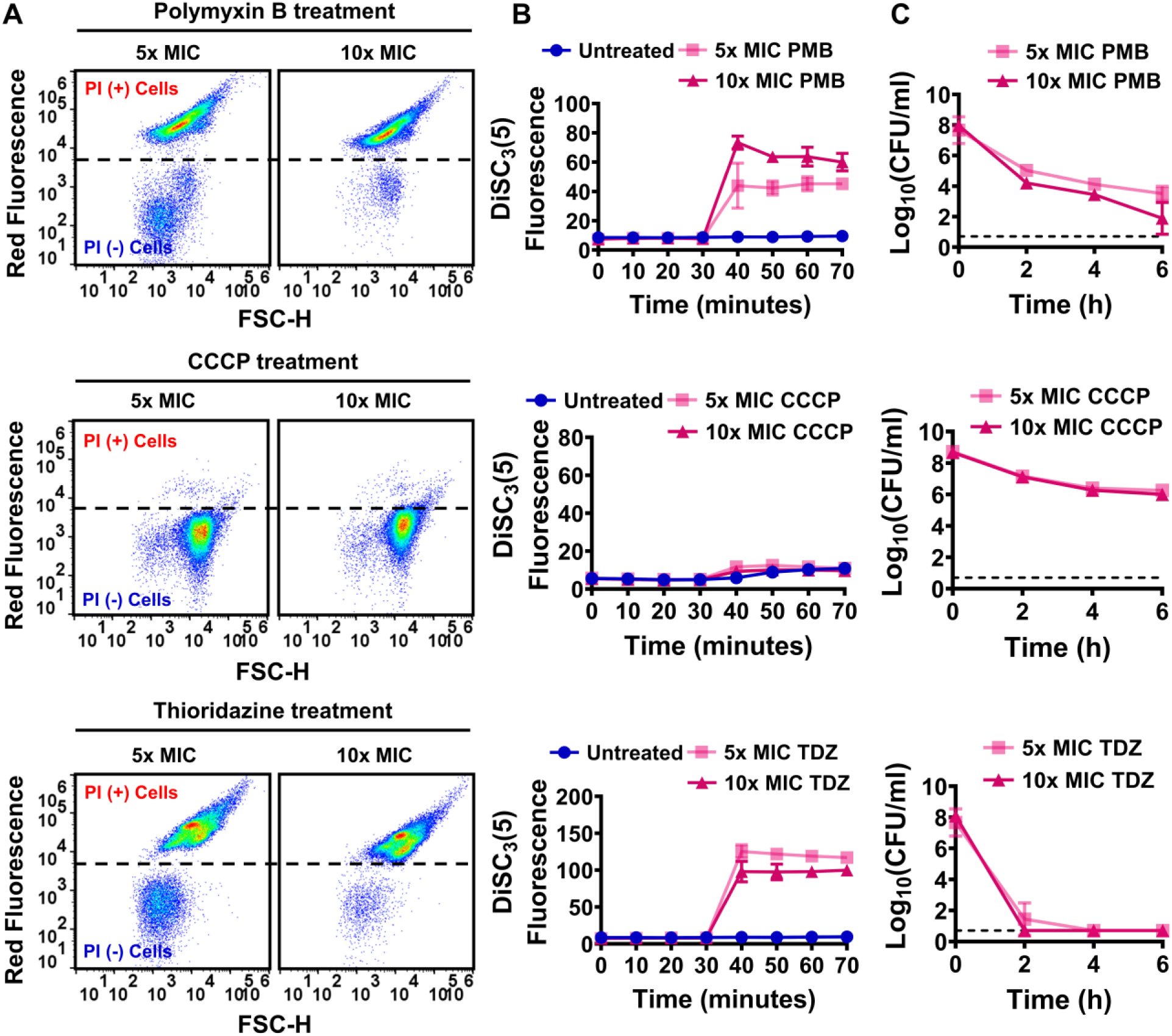
PMF inhibitors increased membrane permeability, disrupted cellular PMF, and reduced persister levels in strain MRSA BAA-41. **(A)** MRSA BAA-41 cells were grown to the exponential phase (OD_600_ of ∼0.1) in Mueller–Hinton broth and treated with polymyxin B (PMB), CCCP, or thioridazine (TDZ) at concentrations of 5× and 10× MIC (**Supplementary Table S1A**). After 1 h treatment, cells were collected and stained with PI (20 μM) dye for flow cytometry analysis. Live and ethanol-treated (70%, v/v) dead cells were used as negative (–) and positive (+) controls **(Supplementary Fig. S2**). A representative flow cytometry diagram is shown here; all independent biological replicates produced similar results. **(B)** Cells grown to the exponential phase (OD_600_ of ∼0.1) were transferred to DiSC_3_(5) assay buffer (50 mM HEPES, 300 mM KCl, and 0.1% glucose) and stained with DiSC_3_(5). When the cells reached an equilibrium state (*t* = 30 minutes), they were treated with polymyxin B, CCCP, or thioridazine at the indicated concentrations. The fluorescence levels were measured with a plate reader at designated time points. Cultures stained with the DiSC_3_(5) but not treated with PMF inhibitors were used as control. **(C)** Cells at the exponential phase (OD_600_ of ∼0.1) were treated with the drugs at the indicated concentrations for 6 h. At designated time points during treatments, cells were collected, washed to remove the chemicals, and spotted on Mueller–Hinton agar plates to obtain CFU counts. Dashed lines in panel C indicate the limit of detection. The number of biological replicates (n) = 3. Data points represent mean ± SD.

**Fig. 2.**
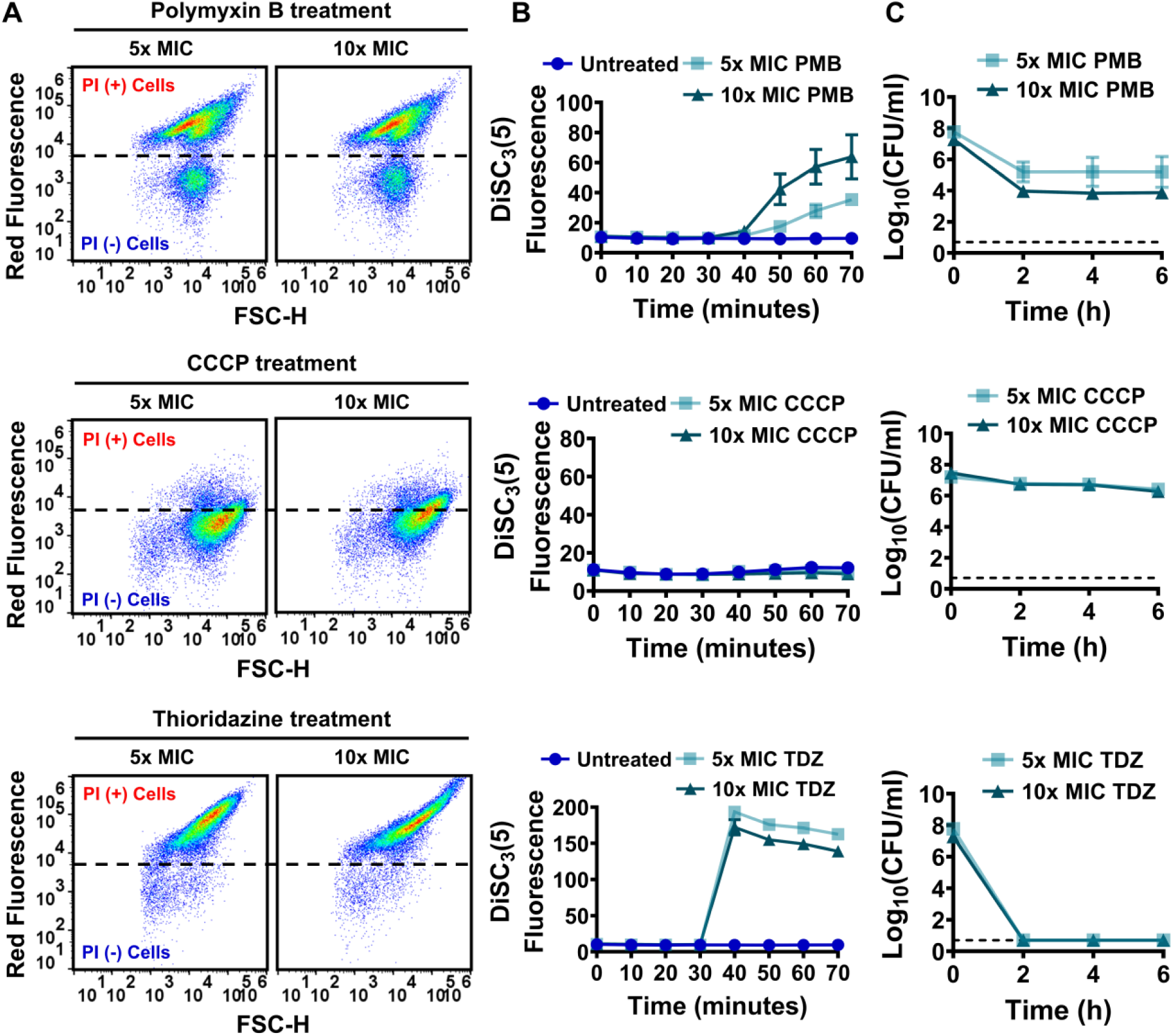
PMF inhibitors increased membrane permeability, disrupted cellular PMF, and reduced persister levels in strain MRSA 700699. Effects of polymyxin B (PMB), CCCP, and thioridazine (TDZ) treatments on cell membranes (**A**), PMF (**B**), and persister levels (**C)** of MRSA 700699 cells were determined as described in **Fig. 1**. A representative flow cytometry diagram is shown here; all independent biological replicates (n = 3) produced similar results. Dashed lines in panel C indicate the limit of detection. Data points represent mean ± SD.

To determine whether the observed membrane permeabilization was linked to the perturbation of PMF, we used the potentiometric probe 3,3′-dipropylthiadicarbocyanine iodide [DiSC_3_(5)] (see **Materials and Methods**), which accumulates on polarized membranes and self-quenches its fluorescence^27,31^. Hyperpolarization due to perturbation of ΔpH enhances the accumulation of DiSC_3_(5) and reduces the fluorescence signals, whereas disruption of ΔΨ increases fluorescence by releasing the probe into the medium^27,31^. Polymyxin B at 5× and 10× MIC disrupted the cellular PMF by selectively dissipating ΔΨ in both strains in a concentration-dependent manner (**Fig. 1B, Fig. 2B**, and **Supplementary Table S2A**). The dissipation of ΔΨ was greater in thioridazine-treated cultures than in polymyxin B–treated cultures (**Fig. 1B, Fig. 2B**, and **Supplementary Table S2A**). Thioridazine at 10× MIC increased the DiSC_3_(5) fluorescence level more than 11-fold in MRSA BAA-41 cells and more than 14-fold in MRSA 700699 cells when compared to untreated controls (**Fig. 1B, Fig. 2B**, and **Supplementary Table S2A**). However, CCCP at 5× and 10× MIC did not disrupt PMF in either strain (**Fig. 1B, Fig. 2B**, and **Supplementary Table S2A**).

We performed clonogenic survival assays to assess the effectiveness of these PMF inhibitors as antipersister drugs. MRSA BAA-41 and MRSA 700699 cells were treated with the inhibitors at 5× and 10× MIC for 6 h to generate kill curves, given that the presence of persisters in a population leads to biphasic kill curves^45^. These assays revealed that CCCP was ineffective against MRSA strains and polymyxin B was unable to eradicate persister cells after 6 h of treatment at the tested concentrations (**Fig. 1C** and **Fig. 2C**). However, thioridazine, which disrupted cell membranes and PMF to a greater extent than the other tested drugs, reduced persister levels of both strains to below the limit of detection at both concentrations tested (**Fig. 1C** and **Fig. 2C**). Although a direct comparison of the effects of these three inhibitors on bacterial cell physiology, including their effects on persistence, might be difficult to obtain due to the concentration-dependent nature of these effects, our data show that the conditions that lead to enhanced membrane permeabilization and PMF disruption may eliminate persister cells.

### Conventional antibiotics do not eliminate MRSA persisters

Next, we investigated whether similar correlations between membrane integrity, PMF levels, and persistence are observed when cells are treated with conventional antibiotics. We selected seven antibiotics, including kanamycin and gentamicin (aminoglycosides that inhibit protein biosynthesis by binding to the 30S ribosomal subunit)^46^, ampicillin (a beta-lactam that inhibits cell wall biosynthesis by binding to penicillin-binding proteins)^47^, ofloxacin and ciprofloxacin (quinolone antibiotics that block DNA synthesis by inhibiting DNA gyrase/topoisomerase)^48^, fosfomycin (a phosphonic acid that blocks cell wall biosynthesis by inhibiting the initial step involving phosphoenolpyruvate synthetase)^49^, and vancomycin (a glycopeptide antibiotic that inhibits cell wall biosynthesis by binding to the growing peptide chain)^50^. Using commercial strips, we confirmed that the MICs of kanamycin, gentamicin, ampicillin, ofloxacin, ciprofloxacin, fosfomycin, and vancomycin for strain MRSA BAA-41 were within the standard test ranges (**Supplementary Table S1B**). As both kanamycin and gentamicin have a similar mode of action, kanamycin was selected for persister assays for this strain. MICs of ampicillin, ofloxacin, ciprofloxacin, and vancomycin were detectable for strain MRSA 700699, but this strain exhibited high resistance to kanamycin, gentamicin, and fosfomycin. We were unable to determine the MICs of these three antibiotics for strain MRSA 700699, which exceeded the standard test ranges (**Supplementary Table S1B**).

Exponential-phase cells (OD_600_ of ∼0.1) (**Supplementary Fig. S1**) of strains MRSA BAA-41 and MRSA 700699 were treated with conventional antibiotics at 5× and 10× MIC (**Supplementary Table S1B**) for PI staining, DiSC_3_(5), and clonogenic survival assays as described above. MRSA BAA-41 was highly tolerant to kanamycin, ofloxacin, and ciprofloxacin, and these antibiotics neither permeabilized the cytoplasmic membrane nor dissipated the PMF of this strain at the concentrations tested (**Supplementary Fig. S3A–C**). Ampicillin, fosfomycin, and vancomycin were able to permeabilize the cell membrane without altering the PMF of strain MRSA BAA-41 but did not eradicate the persister cells of this strain at the concentrations tested (**Supplementary Fig. S3A–C**). Similar trends were observed for strain MRSA 700699 (**Supplementary Fig. S4A– C**). Although ampicillin and vancomycin significantly permeabilized MRSA 700699 cells, at the concentrations tested, none of the antibiotics altered the cellular PMF or eradicated persister cells of this strain (**Supplementary Fig. S4A–C**). Altogether, the results of PMF inhibitor and conventional antibiotic treatments suggest that chemicals that increase both PMF dissipation and membrane permeabilization might be effective antipersister drugs. However, a statistical analysis is necessary to clarify whether PMF dissipation and membrane permeabilization can truly predict persister levels.

### Simple multivariable regression analysis identifies a linear correlation between independent and response variables

The patterns we observed among membrane permeabilization, PMF dissipation, and persister levels after treatment with known PMF inhibitors and conventional antibiotics suggest a correlation between these parameters. When we generated a membrane permeability vs. PMF disruption plot using data from all independent biological replicates for all combinations of MRSA strains and drug concentrations (**Fig. 3A**), we observed two distinct clusters. The first cluster in this two-dimensional plot (**Fig. 3A**, red circle) primarily represents the data points corresponding to conventional antibiotics. Although some of these antibiotics (e.g., ampicillin, fosfomycin, and vancomycin) permeabilized cell membranes, they did not necessarily dissipate cellular PMF, indicating that these two parameters are not always related. The second cluster (**Fig. 3A**, blue circle) comprises the drugs that perturb PMF (e.g., thioridazine and polymyxin B). These drugs drastically permeabilized the cell membranes of both strains independent of PMF disruption and were more effective against persister cells than the drugs in the first cluster (**Fig. 3A**). The data on chemicals in the second cluster may indicate either a lack of correlation between membrane permeabilization and PMF disruption or the existence of a threshold level for PMF disruption that leads to drastic membrane permeabilization. If we assume that PMF dissipation and membrane permeabilization are two independent variables, and the persister outcome is the response variable, then the potential two-way interaction between the independent variables should be statistically verifiable.

**Fig. 3.**
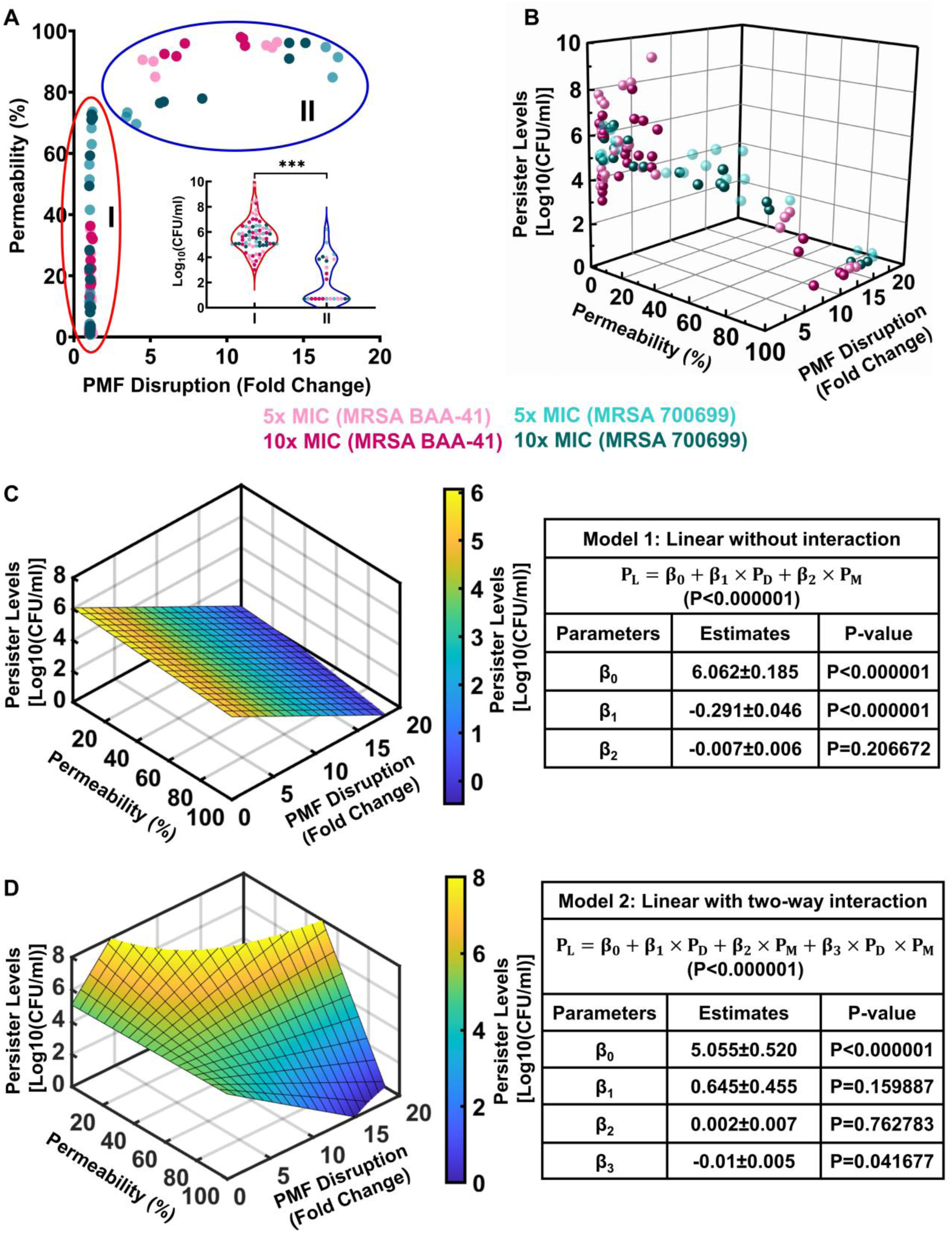
Simple multivariable regression analysis correlates the disruption of PMF and membrane permeability to persister levels. **(A, B)** Two- and three-dimensional scatter plots including all data points for PMF inhibitors and conventional antibiotics for all concentrations and strains tested. In panel A, the red circle indicates cluster I, and the blue circle indicates cluster II. The persister levels corresponding to each cluster are presented in the inset. A Student’s *t*-test with unequal variance was used to find the statistical significance between the persister levels of clusters I and II (***P < 0.0001). (**C**) Multivariable linear regression analysis without an interaction between the independent variables. (**D**) Multivariable linear regression with a two-way interaction between the independent variables. *P*_*L*_ = persister level; *P*_*D*_ = PMF disruption; and *P*_*M*_ = membrane permeabilization. F statistics were used for the statistical analysis with the threshold value set to P = 0.01.

Our three-dimensional scatter plot of membrane permeability, PMF disruption, and persister-level data may indicate a linear correlation between the independent and response variables (**Fig. 3B**). To test whether a two-way interaction exists between the independent variables, we performed a simple multivariable correlation analysis in which the response is predicted by the independent variables using two different linear model equations with or without an interaction term (β_3,_ **Fig. 3C, D**). The first model equation without the interaction term indicates that PMF disruption has a significant effect on persister level (P < 0.0001), but membrane permeability has a comparatively smaller effect (P = 0.2067) (**Fig. 3C, D**). Although the analysis associated with the second model equation may suggest the existence of interaction between the independent variables, the F statistics used to compare the model equations indicate that the first model fits the experimental data better than the second model (P < 0.01) (**Fig. 3C, D**). However, both regression models fit the experimental data better than a model that contains no independent variables (P < 0.00001).

Our experimental data, together with the statistical analysis, demonstrate the importance of cellular PMF dissipation on MRSA persister levels, regardless of the strains used. Although these model equations may not predict the exact number of persister cells, they may predict the conditions necessary to reduce the level of persister cells to below the limit of detection. When we calculated the minimum PMF disruption required to eradicate persister cells if 90% of the cells are assumed to be permeabilized, the first and second model equations revealed that at least 16.22 ± 4.72-fold and 17.85 ± 3.91-fold PMF disruption, respectively, is required to reduce MRSA persister levels to below the limit of detection [5 colony-forming units (CFU)/ml], which is consistent with our experimental data (**Fig. 1** and **Fig. 2**). However, whether PMF inhibitors can truly be used as antipersister drugs requires further validation, as our current analysis includes a limited number of PMF inhibitors.

### High-throughput screening identified new PMF inhibitors for the MRSA strains

To identify additional PMF inhibitors, we screened a small chemical library, MitoPlate I-1, containing 22 mitochondrial inhibitors. Each chemical was tested at four different concentrations in the wells of a 96-well plate. These chemicals included complex I inhibitors (rotenone and pyridaben), complex II inhibitors (malonate and carboxin), complex III inhibitors (antimycin A and myxothiazol), uncouplers [trifluoromethoxy carbonylcyanide phenylhydrazone (FCCP) and 2,4-dinitrophenol], ionophores (valinomycin and calcium chloride), and other chemicals (gossypol, nordihydroguaiaretic acid, polymyxin B, amitriptyline, meclizine, berberine, alexidine, phenformin, diclofenac, celastrol, trifluoperazine, and papaverine) that directly or indirectly inhibit the ETC of mitochondria^51,52,61,62,53–60^. Although this library was specifically designed for mammalian cells, we reasoned that some of the chemicals might be effective for bacteria as the ETC is evolutionarily conserved^63^. Exponential-phase cells (OD_600_ of ∼0.1) of strains MRSA BAA-41 and MRSA 700699 were used to perform the DiSC_3_(5) assay for our initial screening. For both strains, alexidine, diclofenac, celastrol, trifluoperazine, and amitriptyline selectively dissipated ΔΨ [increase in DiSC_3_(5) fluorescence levels compared to untreated control], whereas nordihydroguaiaretic acid and gossypol selectively dissipated ΔpH [decrease in DiSC_3_(5) fluorescence levels compared to untreated control] (**Supplementary Fig. S5** and **Fig. S6**). FCCP and antimycin A particularly disrupted the PMF in strain MRSA BAA-41 (**Supplementary Fig. S5**).

We performed PI staining, DiSC_3_(5), and clonogenic survival assays to verify the reproducibility and efficacy of the identified chemicals against MRSA persisters. Exponential-phase cells (OD_600_ of ∼0.1) of strains MRSA BAA-41 and MRSA 700699 were treated with the identified drugs at 5× and 10× MIC. A two-fold macro-dilution method^64^ was used to determine the MICs of these drugs (**Supplementary Table S1C**). The MIC of antimycin A is much higher than the range we tested (0.0078125–2 mM); therefore, antimycin A was not tested in the persister response assays. Our results showed that nordihydroguaiaretic acid and gossypol drastically perturbed the PMF by dissipating ΔpH, robustly permeabilized cell membranes, and reduced persister levels to below the limit of detection within 6 h of treatment at the concentrations tested for both MRSA BAA-41 and MRSA 700699 (**Fig. 4A–C** and **Fig. 5A–C**). The potency of gossypol in targeting cellular PMF seemed to be quite high, as it reduced DiSC_3_(5) fluorescence levels more than 122-fold at 10× MIC compared to untreated cells (**Fig. 4B, Fig. 5B**, and **Supplementary Table S2C**). Trifluoperazine and amitriptyline similarly reduced persister levels to below the limit of detection for both strains; however, these drugs potentially permeabilized the cell membrane by dissipating ΔΨ (**Fig. 4A–C** and **Fig. 5A–C**). Alexidine, FCCP, diclofenac, and celastrol affected persister levels, cellular PMF, and membrane permeabilization in a concentration-dependent manner for both strains (**Supplementary Fig. S7A–C** and **Fig. S8A–C**). Although conditions that drastically disrupted cellular PMF and permeabilized the membrane (e.g., alexidine treatment at 10× MIC) reduced persister levels to below the limit of detection (**Supplementary Fig. S7A–C, Fig. S8A– C**, and **Table S2C**), conditions that barely perturbed PMF and cell membrane permeabilization (e.g., celastrol treatments at 5× and 10× MIC) were ineffective in eliminating persister cells (**Supplementary Fig. S7A–C, Fig. 8A–C**, and **Table S2C**).

**Fig. 4.**
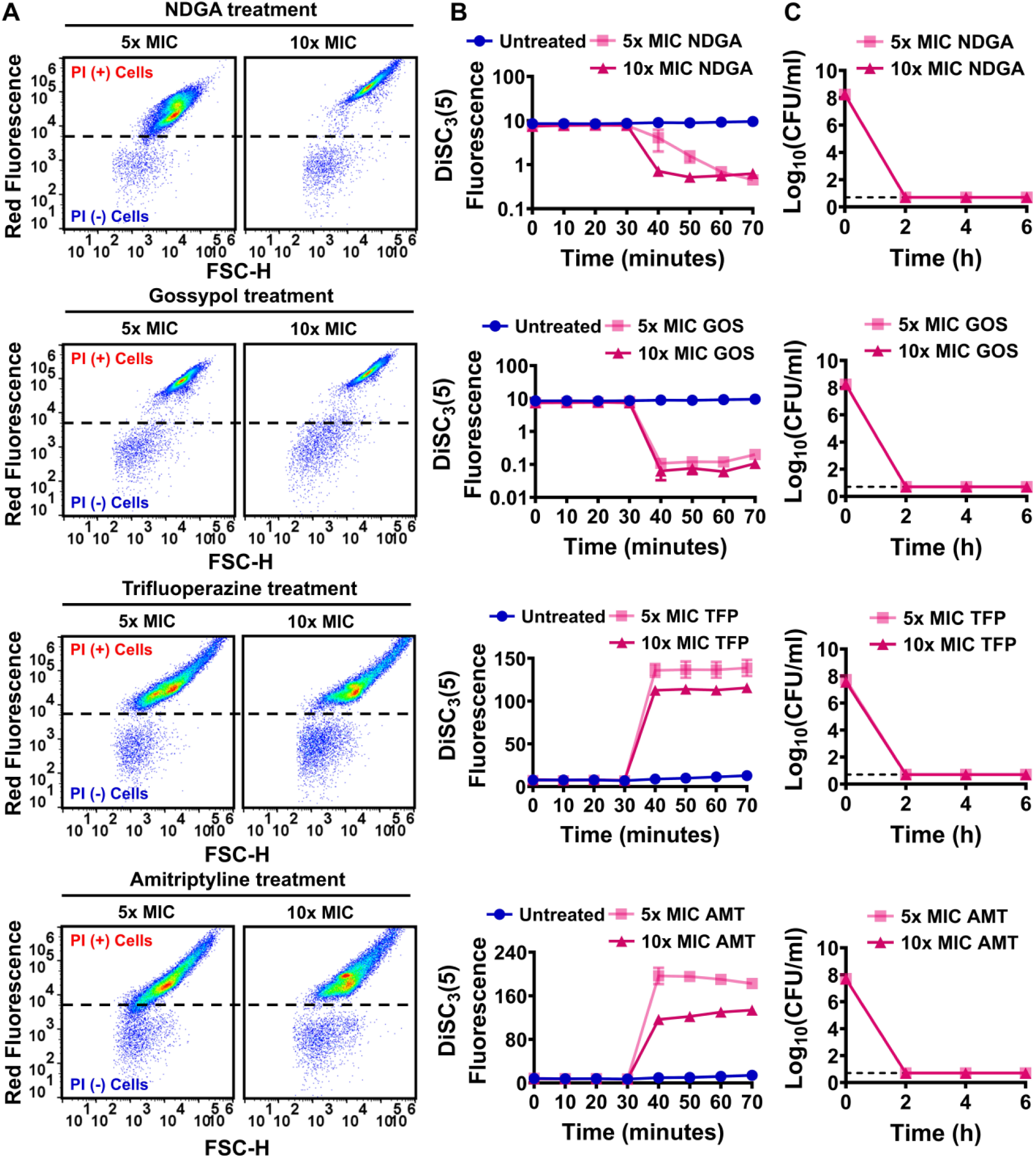
Identified drugs increased membrane permeability, disrupted cellular PMF, and reduced persister levels in strain MRSA BAA-41. Effects of nordihydroguaiaretic acid (NDGA), gossypol (GOS), trifluoperazine (TFP), and amitriptyline (AMT) treatments on cell membranes (**A**), PMF (**B**), and persister levels (**C)** of MRSA BAA-41 cells were determined as described in **Fig. 1**. A representative flow cytometry diagram is shown here; all independent biological replicates (n = 3) produced similar results. Dashed lines in panel C indicate the limit of detection. Data points represent mean ± SD.

**Fig. 5.**
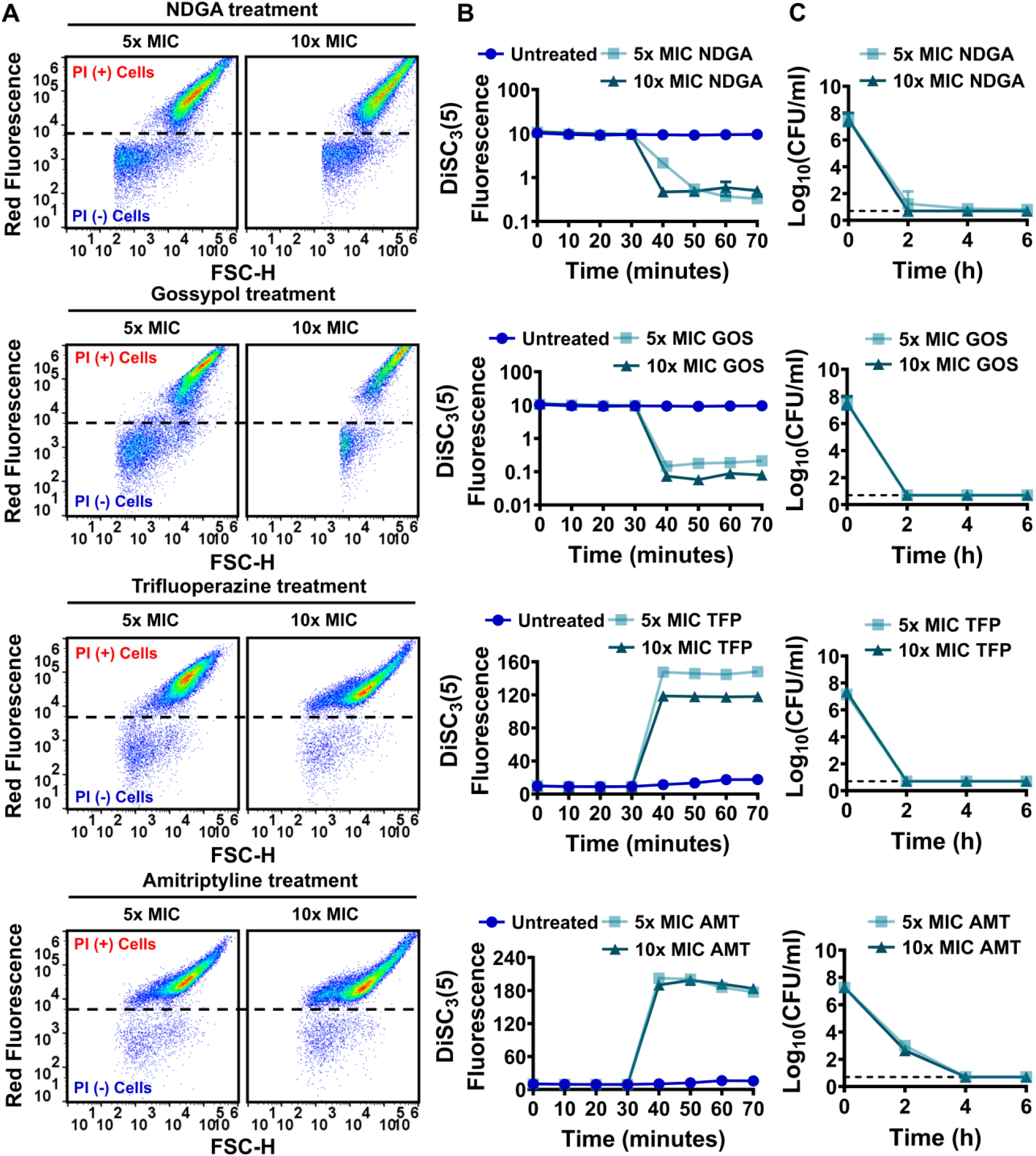
Identified drugs increased membrane permeability, disrupted cellular PMF, and reduced persister levels in strain MRSA 700699. Effects of nordihydroguaiaretic acid (NDGA), gossypol (GOS), trifluoperazine (TFP), and amitriptyline (AMT) treatments on cell membranes (**A**), PMF (**B**), and persister levels (**C)** of MRSA 700699 cells were determined as described in **Fig. 1**. A representative flow cytometry diagram is shown here; all independent biological replicates (n = 3) produced similar results. Dashed lines in panel C indicate the limit of detection. Data points represent mean ± SD.

The results of our screening assay support our initial analysis, highlighted in **Fig. 3**. When we repeated our statistical analysis by combining the new and initial data sets of independent variables (PMF disruption and membrane permeability) for all drugs and conditions, we found that the first model (without a two-way interaction) fit the experimental data better than the second model (P < 0.01) (**Supplementary Fig. S9A, B**). Although both PMF disruption and membrane permeability had significant effects on persister levels (P < 0.0001), the effects of interactions between PMF and membrane permeability on persistence were insignificant with the addition of new data (P = 0.1495). Altogether, our results verified that conditions leading to robust disruption of PMF and drastic cell membrane permeabilization could reduce persister levels to below the limit of detection.

## DISCUSSION

In this study, the strains MRSA BAA-41 and MRSA 700699 were employed to explore the disruption of PMF as a potential therapeutic approach against MRSA persister cells. These strains are *S. aureus* clinical isolates that are intrinsically resistant to methicillin^26,65^. MRSA BAA-41 was isolated from a patient in a New York City hospital in 1994^65^. MRSA 700699 was isolated from the pus and debrided tissue that developed at a surgical incision in the sternum of an infant from Japan^26^. The two strains have different growth rates—MRSA BAA-41 proliferates faster than MRSA 700699 in Mueller–Hinton broth (**Supplementary Fig. S1**)—and are both highly tolerant to conventional antibiotics.

Our initial data sets obtained from known PMF inhibitors and conventional antibiotics highlight a strong correlation between cellular membrane permeabilization, PMF disruption, and persister levels in MRSA strains. Our statistical analysis demonstrated that the two independent variables (membrane permeabilization and PMF disruption) had a significant effect on the response variable (persister levels). We further showed that the response variable can be defined by a linear regression model with an insignificant two-way interaction between the independent variables. However, this lack of statistical interaction does not necessarily imply that PMF and membrane integrity are not related. PMF inhibitors seem to permeabilize cell membranes either completely (e.g., thioridazine) or not at all (e.g., CCCP), depending on their potency; therefore, permeabilization mediated by PMF inhibitors could potentially occur above a certain potency threshold. Because our experimental results and data analysis suggest that PMF inhibitors can be effective antipersister drugs or adjuvants for MRSA strains, we screened a small chemical library containing 22 mitochondrial inhibitors and found that several drugs, including nordihydroguaiaretic acid, gossypol, trifluoperazine, amitriptyline, and alexidine, were effective PMF inhibitors for MRSA strains and could robustly permeabilize the cell membrane and reduce persister levels to below the limit of detection.

The chemicals in the library inhibit different mechanisms of the mitochondrial ETC system^51,52,61,62,66–68,53–60^. The ETC is evolutionarily conserved across species^63^, which may explain the observed high hit rate achieved by screening a small chemical library. As cancer cells are characterized by increased proliferation and mitochondrial activities, these drugs are effective inhibitors for many cancer cells. Gossypol is a naturally occurring aldehyde extracted from a cotton plant that inhibits two fragments of mitochondrial electron transfer and triggers the production of reactive oxygen species^62^, which has antitumor effects against several myeloma cells by inducing apoptosis^69^. Trifluoperazine is an antipsychotic drug that dissipates mitochondrial transmembrane potential, permeabilizes the plasma membrane, and decreases the viability of hepatoma tissue culture cells *in vitro*^68^. Amitriptyline is a tricyclic antidepressant drug that inhibits the activities of mitochondrial complex III and stimulates the generation of reactive oxygen species in human hepatoma cells^70^. Other identified drugs, including nordihydroguaiaretic acid, alexidine, and celastrol, induce mitochondrial apoptosis in cancer cells^55,61,71^.

PMF is crucial for bacterial cell growth and survival under normal and/or stress conditions^44^. As the driving force for ATP synthesis via F_1_F_0_-ATPase^44^, PMF provides the necessary energy for many intracellular processes, forming the Achilles heel of living organisms; therefore, the dissipation of one of its components (ΔΨ or ΔpH) can dismantle the cellular adenylate energy charge and kill bacteria^27^. Several studies have demonstrated the importance of PMF for the elimination of bacterial persisters^23,27,72^. Persister cells can consume specific carbon sources and generate PMF through the oxidative ETC, making them vulnerable to the presence of aminoglycosides^23^. Significant reductions in persister levels are observed when ETC activity, the driving force of the PMF, is genetically and chemically repressed^72^. Starvation-induced antibiotic-tolerant cells can be eradicated by disrupting cellular PMF^73^, emphasizing the importance of PMF as an antimicrobial target.

Our screening assay identified a number of drugs that were highly effective against MRSA persisters. In *Escherichia coli*, trifluoperazine irreversibly inhibits ATP synthase by interacting with the F_0_ and F_1_ subunits^74^. Amitriptyline inhibits the AcrB multidrug efflux pump in *Salmonella typhimurium* and *E. coli* strains^75^ and kills drug-resistant gram-positive and -negative bacteria when used as an antibiotic adjuvant^76^. Nordihydroguaiaretic acid disrupts the cytoplasmic membrane and reduces intracellular ATP levels of *S. aureus*^77^. Alexidine has broad-spectrum activities against *Enterococcus faecalis* biofilm infections and fungal pathogens^78^. However, the exact molecular mechanism of action of alexidine against bacteria has yet to be elucidated. Diclofenac inhibits DNA synthesis in *E. coli* and *S. aureus* and exhibits antibacterial activity^79^. In addition, celastrol treatment makes *B. subtilis* cells elongated and spindle-shaped. Using transmission electron microscopy, celastrol has been shown to damage cell membranes to a certain extent^80^. Altogether, although the bactericidal effects of the identified PMF inhibitors (e.g., nordihydroguaiaretic acid, gossypol, trifluoperazine, amitriptyline, and alexidine) have already been highlighted in the literature, their effects on persister cells, to the best of our knowledge, have not been well characterized.

In *E. coli*, thioridazine was previously shown to selectively dissipate ΔpH by potentially interacting with membrane-bound proteins associated with energy metabolism, such as succinate:quinone oxidoreductase (SdhA, SdhB, SdhC, and SdhD); cytochrome bd-I ubiquinol oxidase (CydX); and NADH:quinone oxidoreductase complexes (NuoJ and NuoF)^81^. However, our current study demonstrated that thioridazine disrupts ΔΨ in gram-positive bacteria, underlining the existence of distinct mechanisms across species. Culture conditions (e.g., inhibitor concentrations and the timing of inhibitor addition); redundant interactions between the inhibitors and cellular components; the existence or absence of an outer membrane; and the thickness of peptidoglycans may affect the cellular responses to treatments. Moreover, we found that lower concentrations (5× MIC) of thioridazine, CCCP, FCCP, trifluoperazine, amitriptyline, diclofenac, and celastrol disrupted cellular PMF more than higher concentrations (10× MIC). These PMF inhibitors disrupt ΔΨ, and we did not observe the same phenomenon for inhibitors that selectively dissipate ΔpH, which warrants further investigation.

The rise of antibiotic tolerance is one of the most critical global public health threats of the 21st century, and bacterial persistence contributes to this problem, as persister variants facilitate the recurrence of chronic infections and the emergence of drug-resistant mutants. Here, we demonstrate that PMF inhibitors can be highly effective bactericidal antibiotics with the potential to eradicate persister cells. Our statistical analysis verified that inhibitors that enhance PMF disruption and cell membrane permeabilization could be potent antipersister drugs. The outcomes of this study also support the use of screening strategies^27^ for the development of novel drugs that selectively target bacterial PMF.

## MATERIALS AND METHODS

### Bacterial strains, chemicals, and culture conditions

The strains MRSA BAA-41 and MRSA 700699 used in this study were obtained from Dr. Kevin W. Garey at the University of Houston^26,65^. Chemicals were purchased from Fisher Scientific (Atlanta, GA), VWR International (Pittsburg, PA), or Sigma Aldrich (St. Louis, MO). MitoPlate I-1 (Catalog# 14104) used for chemical screening (**Supplementary Table S3**) was obtained from Biolog, Inc. (Hayward, CA). The chemical library contained four different concentrations (C_1_, C_2_, C_3_, and C_4_) for each drug. However, these concentrations were not disclosed by the vendor.

Mueller–Hinton broth [2.0 g beef extract powder, 17.5 g acid digest of casein, and 1.5 g soluble starch in 1 L deionized (DI) water] was used to grow the MRSA strains. To enumerate the CFU, Mueller–Hinton agar (2.0 g beef extract powder, 17.5 g acid digest of casein, 1.5 g soluble starch, and 17.0 g agar in 1 L DI water) was used. Treated cells were washed with 1× phosphate-buffered saline (PBS) solution to lower the concentrations of antibiotics and chemicals below their MICs. Conventional antibiotics (kanamycin, ampicillin, ofloxacin, ciprofloxacin, fosfomycin, and vancomycin); known PMF inhibitors (CCCP, polymyxin B, and thioridazine); and the hit chemicals obtained from the screening assay (alexidine, nordihydroguaiaretic acid, FCCP, diclofenac, celastrol, gossypol, trifluoperazine, and amitriptyline) were used at 5× and 10× MIC to treat the MRSA strains. MICs of antibiotics and identified chemicals for the two strains are provided in **Supplementary Table S1A–C**. The ETEST strip method was used to determine the MICs of kanamycin, ampicillin, ofloxacin, ciprofloxacin, fosfomycin, and vancomycin. A two-fold serial dilution (macro-dilution) method was used to detect the MICs of CCCP, polymyxin B, thioridazine, alexidine, nordihydroguaiaretic acid, FCCP, diclofenac, celastrol, gossypol, trifluoperazine, and amitriptyline^64^. The vendor, catalog, and purity information of all chemicals is listed in **Supplementary Table S4**. The solvents and stock solution concentrations of chemicals are tabulated in **Supplementary Table S5**. Chemicals dissolved in DI water were sterilized with 0.2-μm syringe filters. An autoclave was used to sterilize liquid and solid media. Overnight pre-cultures were prepared by inoculating cells from a 25% glycerol cell stock (stored at –80 °C) in a 14-ml round-bottom Falcon test tube containing 2 ml Mueller–Hinton broth and cultured at 37 °C for 24 h in an orbital shaker at 250 revolutions per minute (rpm). Main cultures were prepared by diluting overnight pre-cultures 100-fold into 2 ml fresh Mueller–Hinton medium in 14-ml test tubes. Unless otherwise stated, chemical treatments were performed at the exponential phase (OD_600_ of ∼0.1) for 6 h. The shaker speed and temperature were kept constant (250 rpm and 37 °C) in all experiments.

### Cell growth and persister quantitation by clonogenic survival assays

Overnight pre-cultures were diluted 100-fold in 14-ml test tubes containing 2 ml Mueller–Hinton medium and grown in the shaker. At indicated time points, cell samples were collected to measure OD_600_ with a Varioskan LUX Multimode Microplate Reader (Thermo Fisher, Waltham, MA, USA). When the cultures reached an OD_600_ of 0.1, cells were treated with antibiotics or chemicals at 5× and 10× MIC. At designated time points, 200 μl treated cultures were collected and diluted in 800 μl sterile PBS. Diluted cell cultures were then washed twice with PBS by centrifugation at 13,300 rpm (17,000 × *g*) for 3 minutes to remove the antibiotics and chemicals, as described elsewhere^82^. After the final centrifugation, 900 μl supernatant was removed, and the pelleted cells were resuspended the remaining 100 μl, which was then used for a 10-fold serial dilution in 90 μl PBS. Ten microliters of diluted cells were then spotted on Mueller–Hinton agar. Ninety microliters of undiluted cell suspension were also plated on Mueller–Hinton agar to increase the limit of detection (which is equivalent to ∼5 CFU/ml). After incubation of the agar plates for 16 h at 37 °C, CFUs were counted to determine the persister levels. Incubations longer than 16 h did not increase the CFU levels.

### DiSC_3_(5) assay

Overnight pre-cultures were diluted 100-fold in 14-ml test tubes containing 2 ml fresh Mueller– Hinton broth and grown at 37 °C with shaking (250 rpm). Exponential-phase cells (OD_600_ of ∼0.1) were collected, washed three times with a buffer solution (50 mM HEPES, 300 mM KCl, and 0.1% glucose), and centrifuged at 13,300 rpm^31^. After the final washing step, pelleted cells were resuspended in 2 ml buffer, loaded with 1 μM DiSC_3_(5) dye, and incubated in the dark. The fluorescence levels were measured with a plate reader at 620-nm excitation and 670-nm emission wavelengths every 10 minutes. When the fluorescence levels reached an equilibrium state (after 30 minutes), stained cells were treated with chemicals at indicated concentrations and incubated in the dark. At designated time points, 200 μl cells were collected to measure the fluorescence levels. Cultures that did not receive any chemical treatment served as control.

### Chemical screening assay

Overnight pre-cultures were diluted 100-fold in 14-ml test tubes containing 2 ml fresh Mueller– Hinton broth and grown at 37 °C with shaking (250 rpm). Cells at an OD_600_ of ∼0.1 were collected and washed three times in buffer (50 mM HEPES, 300 mM KCl, and 0.1% glucose) with centrifugation at 13,300 rpm. After the final washing step, pelleted cells were resuspended in buffer, loaded with 1 μM DiSC_3_(5) dye, and incubated in the dark. Once the fluorescence levels reached a steady-state (after 30 minutes), 100 μl stained cells were transferred to each well of the MitoPlate I-1 preloaded with chemicals (**Supplementary Table S3**) and incubated in the dark. The fluorescence level of each well was measured with the plate reader at designated time points. Wells without chemicals (A1–A8) served as controls.

### PI staining

Overnight pre-cultures were diluted 100-fold in 14-ml test tubes containing 2 ml fresh Mueller– Hinton broth and grown at 37 °C with shaking. Cells at an OD_600_ of ∼0.1 were treated with the chemicals at indicated concentrations for 1 h. Treated cells were then collected and diluted in 0.85% NaCl solution in flow cytometry tubes (5-ml round-bottom Falcon tubes) to obtain a final cell density of ∼10^6^ cells/ml. The resulting cell suspensions were stained with 20 μM PI dye and incubated at 37 °C in the dark for 15 minutes. Stained cells were collected and analyzed with a flow cytometer (NovoCyte Flow Cytometer, NovoCyte 3000RYB, ACEA Biosciences Inc., San Diego, CA, US). Ethanol (70% v/v)-treated cells (i.e., dead cells) were used as a positive control (PI-positive cells), and PI-stained live cells (PI-negative cells) served as a negative control (**Supplementary Fig. S2A, B**). Forward and side scatter parameters obtained from the untreated live cells were used to gate the cell populations on the flow cytometry diagram^83^. For the fluorescence measurement, cells were excited at a 561-nm wavelength and detected with a 615/20-nm bandpass filter.

### Multivariable linear regression analysis

Multivariable linear regression analysis was performed to determine correlations between the response (persister levels) and independent variables (PMF disruption and membrane permeability). CFU/ml, PMF, and membrane permeabilization data sets used here correspond to the last time points of the related assays. Log-transformed values of CFU/ml obtained from clonogenic survival assays were used to measure persister levels. PMF disruption was defined as the fold change in DiSC_3_(5) fluorescence levels between treated and untreated cells, and membrane permeability was defined as the percentage of PI-positive cells in the flow cytometry diagram. GraphPad Prism 9.3.0 was used to perform the multiple linear regression analysis. The linear model equations without and with a two-way interaction are as follows, respectively:

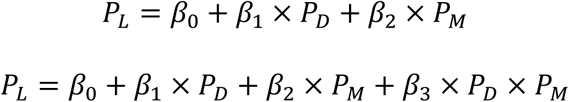

In these equations, *P*_*L*_ is the log-transformed value of the persister levels, *P*_*D*_ is PMF disruption, *P*_*M*_ is membrane permeability, *β*_0_ is the estimate of the model intercept, *β*_1_ is the estimate of the model coefficient of PMF disruption, *β*_2_ is the estimate of the model coefficient of membrane permeability, and *β*_3_ is the estimate of the model coefficient of the interaction term. The parameters identified from the regression analysis were used to generate three-dimensional plots with MATLAB. Quantile–quantile (QQ) probability plots were generated to check the normality of the data set (**Supplementary Fig. S10**).

### Data analysis

Unless stated otherwise, at least three independent biological replicates were performed for each experiment. FlowJo (version 10.8.1) software was used to analyze the flow cytometry data. Each data point in the figures denotes the mean value, and error bars represent the standard deviation (SD). F statistics were used to determine significant differences between the model equations. Student’s *t*-tests with unequal variance were performed to determine the statistical significance between two groups. P-value thresholds were selected as *P < 0.01, **P < 0.001, ***P < 0.0001; ns indicates not significant.

## ACKNOWLEDGMENTS

The authors would like to thank Orman Lab members for their help. This study was supported by an NIH/NIAID R01 AI143643 Award and a University of Houston start-up grant.

## AUTHOR CONTRIBUTIONS

S.G.M., S.G., P.K. and M.A.O. conceived and designed the study. S.G.M., S.G., and P.K. performed the experiments. S.G.M., S.G., and M.A.O. analyzed the data and wrote the paper. All authors have read and approved the manuscript.

## NOTES

The authors declare no competing interests.

